# Host-symbiont-gene phylogenetic reconciliation

**DOI:** 10.1101/2022.07.01.498457

**Authors:** Hugo Menet, Alexia Nguyen Trung, Vincent Daubin, Eric Tannier

## Abstract

**Motivation:** Biological systems are made of entities organized at different scales (*e.g.* macro-organisms, symbionts, genes…) which evolve in interaction. These interactions range from independence or conflict to cooperation and coevolution, which results in them having a common history. The evolution of such systems is approached by phylogenetic reconciliation, which describes the common patterns of diversification between two different levels, *e.g.* genes and species, or hosts and symbionts for example. The limit to two levels hides the multi-level inter-dependencies that characterize complex systems.

**Results:** We present a probabilistic model of evolution of three nested levels of organization which can account for the codivergence of hosts, symbionts and their genes. This model allows gene transfer as well as host switch, gene duplication as well as symbiont diversification inside a host, gene or symbiont loss. It handles the possibility of ghost lineages as well as temporary free-living symbionts.

Given three phylogenetic trees, we devise a Monte Carlo algorithm which samples evolutionary scenarios of symbionts and genes according to an approximation of their likelihood in the model. We evaluate the capacity of our method on simulated data, notably its capacity to infer horizontal gene transfers, and its ability to detect host-symbiont co-evolution by comparing host/symbiont/gene and symbiont/gene models based on their estimated likelihoods. Then we show in a aphid enterobacter system that some reliable transfers detected by our method, are invisible to classic 2-level reconciliation. We finally evaluate different hypotheses on human population histories in the light of their coevolving *Helicobacter pylori* symbionts, reconciled together with their genes.

**Availability:** Implementation is available on GitHub https://github.com/hmenet/TALE. Data are available on Zenodo https://doi.org/10.5281/zenodo.7667342.

## 1 Introduction

The toolbox of evolutionary biology largely relies on the assumption of statistical independence of biological objects at any level of organization: organisms from different species are isolated from a biological system based on their genomes, genomes are cut into independent genes, and inside genes, nucleotides are evolving independently from each other [19].

Yet the essence of living systems lies in dependence: constraint, cooperation or conflict [49]. Symbiotic micro-organisms coevolve with animals or plants [51]. The ensemble they form is gathered under the holobiont concept. It allows to see genes as entities not only following their own interest, not only participating to the functioning of the genome they are hosted by, but also participating to, and probably evolving with, a larger biological system.

A powerful tool to study these inter-dependencies is phylogenetic reconciliation: an ensemble of models and methods explaining the differences and similarities between phylogenies of two entities diversifying concommitantly in evolution. Gene/species systems have been studied by phylogenetic reconciliation, accounting for events of gene duplication, horizontal gene transfer and gene loss (DTL model) [15, 39, 57, 9, 33]. The same model can be applied with little to no modification to symbiont/host [10, 48, 14], protein domain/gene cophylogeny [43, 52], and biogeography has been imagined as one possible level [29, 46, 47]. DTL models have also been used to reconstruct genome histories [17], detect highways of lateral gene transfers in bacteria, archaea or eukaryota [7], assess the relative role of duplication and gene transfer in the evolution of genomes [50], infer ancient symbiotic relationships [5], reconstruct histories of gene fusion and fission [16], model endosymbiotic gene transfer [4].

A limitation of reconciliation methods is their separate application on molecular studies on one side (gene/species cophylogeny), and ecological studies on the other (host/symbiont cophylogeny). The striking methodological unity of the two (the same DTL model is applied on both the molecular and ecological systems) and the growing interest for multi-level systems integrating molecular and ecological inter-dependencies (e.g. the holobiont concept) calls for a unique model for host, symbiont, gene cophylogeny. In support of this claim, a number of empirical studies already rely on host symbiont histories when proposing horizontal gene transfers between symbionts [42, 40, 28, 38], when often, only symbiont gene/species comparisons do not provide enough statistical support for them [61, 44].

Three level reconciliations have been introduced by Stolzer et al. [52] and applied to protein domain, gene and species. They describe two embedded DTL models and an inference method by parsimony. The inference method first reconciles genes and species trees in a DTL model. Then, knowing which genes are present in which species, it reconciles the protein domains with the genes. This defines two kinds of horizontal protein domain transfers between genes, depending on whether the genes are in the same species (which we will call “intra” transfer) or not (”inter” transfer), with a different cost for those two events. Further efforts in this direction have been published by Li and Bansal [26] with a duplication/loss model between gene and species and a DTL model, forbidding inter species transfers, between protein domains and genes. They show NP-hardness of inferring the most parsimonious couple of nested reconciliations [26] and propose different heuristics and problem variants [27, 25]. A probabilistic model without transfers has been proposed by Muhammad, Sennblad, and Lagergren [36]. It aims at inferring dated gene trees from protein domain alignments using Markov Chain Monte Carlo. These attempts prove that it is possible to jointly handle three nested levels in a single computational model. However none of them can yet handle host/symbiont/gene systems in a statistical framework because of specific limitations of each of them: parsimony framework, no transfer or no inter-host transfer, no joint inference between levels of organization, no explicit handling of absent lineages.

We propose a probabilistic model that describes the evolution of three nested entities at three different scales, adapted to a host/symbiont/gene system. In our model a symbiont tree is generated by a DTL model inside the host, with a possibility of evolving temporarily outside the host phylogeny. A gene is generated by a DTL model inside the symbiont, where gene transfer is more likely between symbionts that share a common host (”intra” transfer) than for those that do not (”inter” transfer).

Based on this model we propose an inference method extending the two-level reconciliation “ALE” software [56, 55]. It takes three trees as input, constructs joint scenarios and estimates event rates and likelihoods according to the model. Our implementation also features the possibility to infer a symbiont species tree if only the host tree and several symbiont gene trees are given as input. In addition a comparison of the likelihood of two-level and three-level reconciliations can be used as a test for multi-scale coevolution.

We report a benchmark test of the inference method on simulated data, using an external simulator [24], showing that under the hypothesis that gene transfers are more likely between symbionts of a same host, the three-level reconciliation represent a significant gain compared to the two-level one in terms of the capacity to retrieve the symbiont donors and receivers of horizontal gene transfers.

We use the inference method to identify horizontal gene transfers between *Cinara* aphid symbionts that are detected by expertise [28] but missed by two-level reconciliations.

Finally we show on genes of *Helicobacter pylori* from human populations how likelihood computations can be used to compare different hypotheses on the diversification of a host, given the genes of its symbionts, taking into account the evolutionary dependencies between all three scales.

## 2 2-level reconciliation, definitions and preliminaries

Because we base our model on the two-level DTL reconciliation model implemented in “ALE undated” [55, 34], together with the inference methods, this section is devoted to their brief description.

### 2.1 Model and parameters

We denote by *G, S* respectively a set of gene trees and a species tree, and *δ_D_^S^, δ_S_^T^, δ_S_^L^* is the set of rates at which a gene evolving in a branch of *S* undergo the D,T,L (speciation, duplication, transfer, loss) events. These rates are constant along the species tree and for all gene trees.

The model is generates a rooted phylogenetic tree *G*, given *S* and the rates, according to a birth and death like model. A gene tree can originate in any branch of the species tree with a uniform prior. Speciation occurs at all nodes of *S*, while duplications, transfers and loss can occur along the branches of *S* with the given rates. When a transfer occurs, the receiver branch is chosen according to a uniform probability, avoiding ancestor branches of the donor. This avoids certain impossible transfers but is not sufficient to guarantee that the overall scenario is time feasible. Indeed, two transfers might be incompatible with respect to time [13].

### 2.2 Inference

The core of the inference method consists in computing the probability *P_θ__S_* (*G|S*) of generating *G* given *S* and *θ_S_* = (*p_S_^S^, p_S_^D^, p_S_^T^, p_S_^L^*), the probabilities of S,D,T,L events, proportional to the rates and satisfying *p^S^* + *p^D^* + *p^T^* + *p^L^* = 1. That is, *P_θ_* (*G|S*) is the likelihood of *G*, *S* and *θ_S_*. *S* is assumed binary and rooted. *G* is binary but can be rooted or not. A mapping of the leaves of *G* to the leaves of *S* is needed (the species in which each extant gene is found).

We call *reconciliation scenario* a list of events of kind D, T, L, or S associated to each internal gene tree node, that can be the result of the birth and death process.

These lists transcribe into a mapping of the nodes of *G* to the nodes of *S*. We note *R_G,S_* the set of all possible reconciliation scenarios by which *G* can be produced from *S*. The likelihood of a scenario *r ∈ R_G,S_*, *P_θ__S_* (*r|S*) is the product of the probabilities of all events. Thus we have

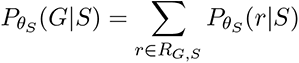

We do not need to fully enumerate all scenarios to compute this sum. Indeed, a dynamic programming scheme along *S* and *G* allows us to sum over scenarios individually on each branch of *S* and ensures tractability. The dynamic programming scheme consists first in a “forward step” traversing the nodes of *G* and *S* in post-order: a node is examined only if its children have been examined before.

If *e, u* are nodes of *S* and *G*, *f* and *g* are descendants of *e*, *v* and *w* are descendants of *u* (if any of these do not exist the corresponding terms must be dropped), and *P_e,u_*= *P_θS_* (*e, u*) is the probability of generating the subtree of *G* rooted at *u* in the subtree of *S* rooted at *e*, then:

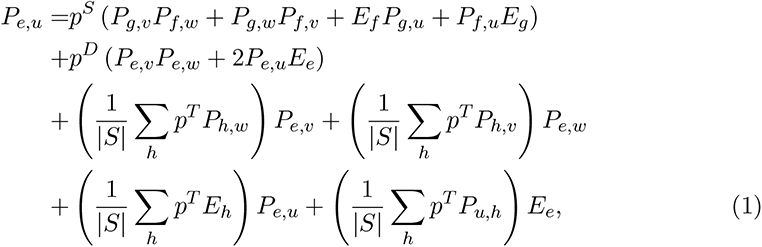

where *E_e_* is the probability that a gene on branch *e* of *S* goes extinct:

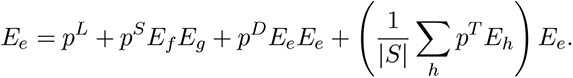

The sum of probabilities at the root of *G*, for all node of *S*, gives *P_θS_* (*G|S*).

A “backward step” then traverses the nodes in the reverse order. It allows one to sample the scenarios based on their probability, or to select the scenario that maximises the marginal likelihood [62]: this means, at each step of the backtracking procedure we select the scenario with maximum likelihood.

Note that this is different from finding the most likely reconciliation scenario. It is possible to find it by a similar procedure, storing the maximum probability in the forward step instead of the sum of the probabilities, and computing the scenario realizing this maximum in the backward step, as in a parsimony algorithm. We did not use this possibility, sticking to the ALE principle.

### 2.3 About time consistency

Simulated scenarios according to the model, and inferred reconciliations do not need to be *time consistent*: a set of transfers might indicate histories that are not feasible on a timeline. This is a known drawback of undated models [13]. There have been attempts to investigate this aspects in several directions. For example, Eucalypt [14] or Notung [52] propose to infer only time feasible scenarios, without any guarantee that such a scenario exists or is can be found in reasonable computing time. Producing a time feasible scenario is NP-complete. Moreover, inferring only time feasible scenarios for one gene family or one symbiont, depending on the biological context, does not guarantee that the combination of scenarios from several gene families or several symbiont will be time consistent: a set of transfers from different genes might not be consistent.

Producing time consistent scenarios with several gene families or symbiont goes back to producing a dated tree [11].

On the other hand, measuring the degree of inconsistency can make this hindsight a strength: for example, one can compare scenarios in relation to this consistency, with the assumption that the more consistent the scenario, the more realistic it is. We used this to compare 2-level and 3-level scenarios in the evaluation of the method by simulations.

## 3 3-level reconciliation, likelihood estimation and scenario inference

### 3.1 Elements of the probabilistic model

The 3-level model is based on two nested 2-level models based on the one presented in the previous section, with the following extensions and restrictions.

#### 3.1.1 Host/Symbiont

A *host* tree *H* is unique, given, rooted. Inside *H*, a *symbiont* tree *S* is generated with the DTL 2-level model, with parameters *δ_H_^D^, δ_H_^L^, δ_H_^T^*, adding the possibility for a symbiont to live temporarily in an unknown host.

Indeed, in the course of their evolutionary history, some symbionts may live outside a host, or within an unknown host. This is a general interesting feature, and is particularly important for us because we invoke unknown hosts in the inference process in the case of inter host horizontal gene transfers (section 3.3).

The utility of this model addition is visible in the *Cinara* aphids example developed in the Results section (see Fig. 6).

#### 3.1.2 Host/Symbiont/Gene

The evolution of a gene tree *G* inside *H/S* reconciliations also follows an adaptation of the DTL model. *G* is generated in one or several symbiont trees with duplication, loss and intra horizontal transfer, with rates *δ_S_^D^, δ_S_^L^, δ_S_^T^*. “Intra” means that horizontal transfer is possible only between symbiont branches (from the same symbiont tree or not) that are present in the same host branch (as the trees *S* are generated in *H*).

Note that the gene tree *G* can refer to a family of genes found in symbionts as well as the host. In the latter case, to remain generic, we simply assume that *S* can be a copy of *H*, reconciled with *H* with only speciations. That is, the host genes are contained in a specific compartment and can transfer to a symbiont, and be transferred from a symbiont.

An illustration of the realization of such a model is given in Figure 1.

**Figure 1:**
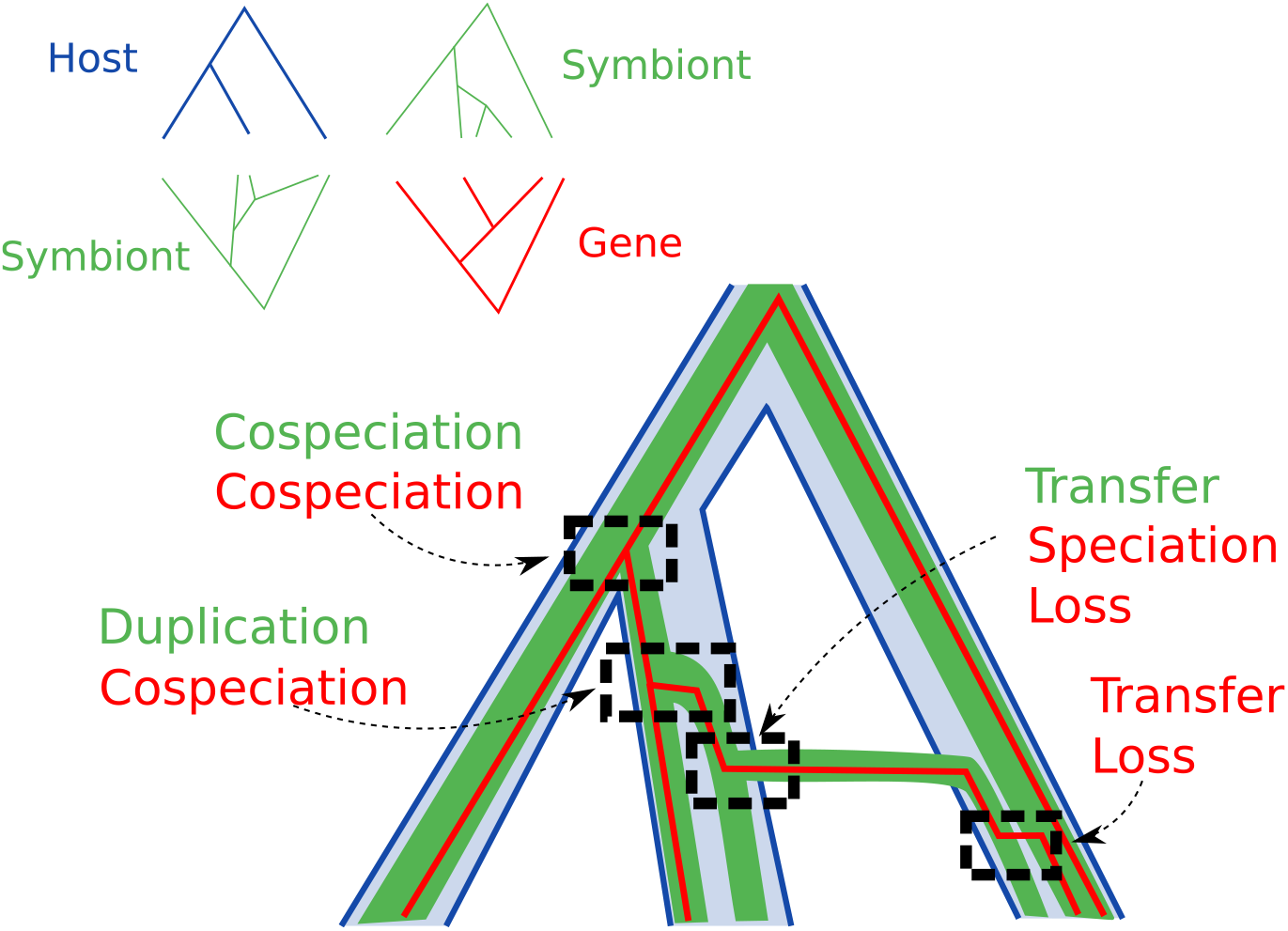
An example of a 3-Level reconciliation input (top left, with three trees and associations between the leaves of two couples of trees) and a possible reconciliation scenario for this input. Events of the host/symbiont co-evolution are written in red, while events of the symbiont/gene reconciliation are written in green.

This model can be immediately used for simulations, but we chose to use an external simulator for our tests [24]. Though this does not allow an identifiability study, which we postpone to a future work, it controls some of the effects of similarities in models and implementation between simulation and inference, providing more difficult instances for testing.

### 3.2 Monte Carlo approximation of the likelihood

Like in the previous section, the inference consists in estimating the parameters (trees and evolutionary rates) and sampling reconciliation scenarios. We consider as input a single rooted binary tree *H*, one or several rooted or unrooted binary symbiont trees *S* = *{S_i_}*, and one or several rooted or unrooted binary gene trees *G* = *{G_i_}*. Both parameter estimation and sampling are accomplished through a calculation of the probability *P* (*G|S, H*) that gene trees *G* have been generated by the model, given *H*, *S*, and given the DTL probabilities for the two reconciliation levels *θ_S_* = (*p_S_^S^, p_S_^D^, p_S_^T^, p_S_^L^*) and *θ_H_* = (*p_H_^S^, p_H_^D^, p_H_^T^, p_H_^L^*) derived from the rates: the DTL probabilities are proportional to the rates and the sum of all three probabilities is 1.

The probability *P* (*G|S, H*) can be decomposed by summing over all possible host/symbiont reconciliation scenarios *r_S,H_ ∈ R_S,H_*:

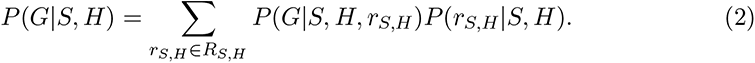

The number of reconciliations in this sum is at least exponential in the size of the input (even the number of scenarios maximizing *P* (*r_S,H_|S, H*) can be exponential [14]). The similar computation in a parsimonious framework is NP-hard [26], so it is probably not possible to exactly and quickly compute *P* (*G|S, H*).

So we apply a Monte Carlo approximation technique. The goal is to sample a reasonable number *N* of symbiont/host reconciliations and approximate *P* (*G|S, H*):

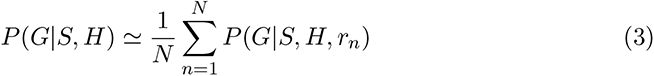

where *r_n_* is sampled in the set *R_S,H_* of all reconciliations according to its likelihood *P* (*r_n_|S, H*). In consequence the term in equation 3 approximates the term in equation 2 according to the Monte Carlo principle.

### 3.3 Reconciliation inference

The computation of *P* (*G|S, H*), as well as sampling reconciliations in *R_S,H_*, is done by successive steps of dynamic programming as shown in Algorithm 1. Steps 2 and 8 are the exact executions of the algorithm ALE [56], with the additional possibility that a symbiont is free living. Free living symbiont are handled by adding a copy of the symbiont tree as an additional host tree. Indeed the reconciliation algorithm can accommodate multiple host trees on separate sets of leaves. Symbiont leaves with no host are matched to themselves instead of a host. In that way, we hypothesize that transfer between free living is less likely than when a common host is known.

Given *r_n_ ∈ R_S,H_*, the probability *P* (*G|S, H, r_n_*) can be computed with an adaptation of the same dynamic programming algorithm (step 15 of Algorithm 1). The only modification is that during the dynamic programming process, for all gene transfer possibilities, it is checked if the donor and receiver symbiont share a host in *r_n_*. If they do, then it is an “intra” transfer and the transfer has the probability defined by the transfer rate.

### 3.4 Inter species transfer through ghost species

Transfer between two symbionts in different hosts is possible through ghost species. Indeed it is always reasonable to assume that a major part of species are extinct or not sampled and gene transfers are often “from the dead” [53, 20, 63, 58].

In consequence, a transfer can have occured from a donor that is now extinct. Figure 2 shows how an “inter” transfer between symbionts *i* and *j* (on the left) can occur, even if it is not explicitly modeled, through a sister lineage to *i*, that switched host and transferred a gene to *j* (on the right). The sister lineage then goes extinct, which explains that the gene looks like it is transferred from *i* to *j*.

**Figure 2:**
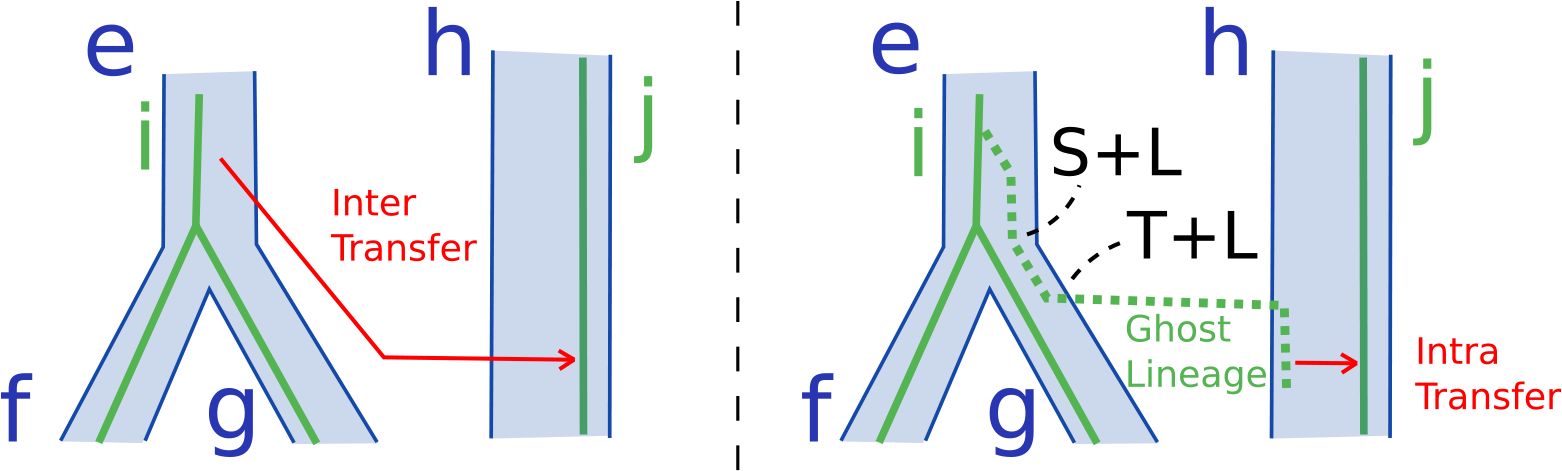
A gene transfer between two symbiont lineages that are in different host lineages (inter transfer) is explained with intra-transfers and ghost lineages. Left part shows the host phylogeny (blue pipes), the reconciled symbiont phylogeny (green lines) and a gene transfer (in red) from lineage *i* to lineage *j*, while *i* is in host lineage *e* and *j* is in host lineage *h*. This direct inter transfer is forbidden by the model. Right part shows a mechanism allowed by the model that has the exact same result, and the way to compute the associated probability. First the symbiont lineage *i* undergoes a speciation and a loss (S+L), and then a transfer and a loss (T+L) before the extinction of the symbiont (or its absence in the taxon sampling) inside *j*. Now the gene transfer (in red) is an intra transfer, as it is transfered between two symbionts inside *h*.

We denote by *P_S_^T^* (*i → j*) the probability for a gene present in symbiont *i* to undergo a horizontal transfer to symbiont *j*, and *P_H_^T^*(*e → h*) the probability for a gene present in a symbiont associated to host *e* to transfer to a symbiont associated to host *h*. Let *H_i_* (*H_j_*) be the set of host branches that contain symbiont *i* (resp. *j*). We go from *P_H_^T^* to *P_S_^T^* by summing over all possibles hosts *h* of the receiver symbiont *j* and all hosts *e* of the donor symbiont *i*:

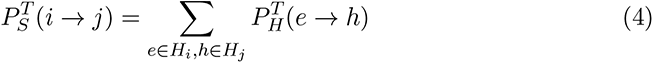

At fixed *h* we rewrite with *P_e_* = *P^T^* (*e → h*). Recall *p_H_^T^* are the probability of horizontal transfer in the symbiont/gene reconciliation, and *p_H_^S^, p_H_^D^, p_H_^T^, p_H_^L^* the probabilities of speciation, duplication, transfer and loss in the host/symbiont reconciliation.

Let *E_e_* be the probability of extinction, that is, the probability that a gene is present in a branch *e* of the host tree and absent from all the leaves. Let *|S_h_|* be the number

**Algorithm 1.**
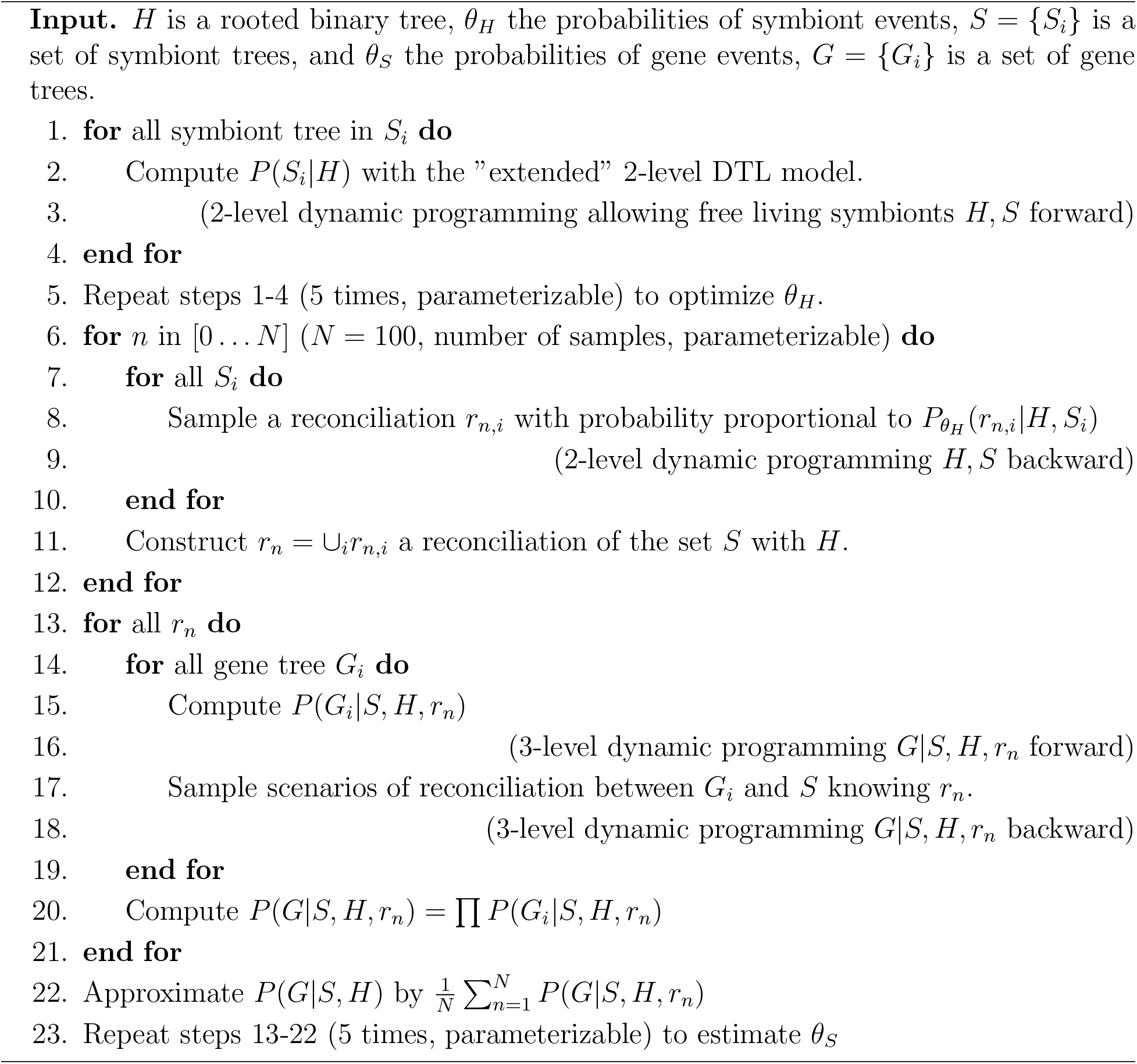
The Monte Carlo inference algorithm.

of symbiont branches matched to host *h* in the host/symbiont reconciliation scenario. The initial case in our inductive definition of *P_e_* = *P^T^* (*e → h*) is the case where *e* = *h*, so when the donor symbiont is in one of the receiver symbiont host, in that case the probability to transfer to that one symbiont of *h*, is uniform among the *|S_h_|* symbionts present in *h*. Then, for the induction, we rewrite the undated reconciliation equations, to progress a symbiont in the host tree from any host *e* to host *h* of the receiver symbiont and such that the symbiont species we invoke then goes extinct. The notations are similar to those used in the undated ALE description in [34], Section 2 or figure 2: we denote by *f, g* the children of a host *e*, and by *|H|* the number of nodes in *|H|*.

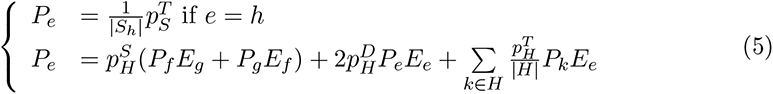

Note that the last sum in the equation is limited to the *k* that are not ancestors of *e*, as in ALE. This equation has a self dependency due to the Transfer/Loss event, which is already accounted for in reconciliation methods [21, 56]. We forbid successions of several Transfer/Loss events to break this self dependency and solve this equation.

### 3.5 Sequential and 2-level estimation of the likelihood

Because the Monte Carlo approach can be computationally heavy, we devised an alternative “Sequential” heuristic. Instead of sampling scenarios randomly like in the Monte Carlo, we select only one of them, maximizing the marginal likelihood [62].

That is, at each step of the backtracking of the dynamic programming procedure we select the event maximizing the probability in the sum of Equation (1). In other words, we decompose *P*_(_*_θS,θH_* _)_(*G|S, H*) into

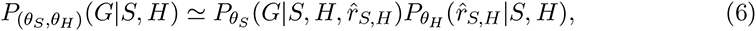

where *r̂_S,H_* is the reconciliation scenario maximizing the marginal likelihood. Note that is can be different from taking the most likely scenario, which is also a possible strategy, consisting in changing the Equation (1) from a sum to a max, and backtracking in this alternative dynamic programming table. So this variant consists in removing the “for” loop of step 6 of Algorithm 1 and replacing step 8 by a systematic choice of a maximum instead of choosing in the sum of Equation 1 with probabilities proportional to the term values.

This approach is similar to the one of Stolzer et al. [52]. The differences are, apart from using a probabilistic setting, that we use marginal likelihood, and that we compute the inter transfer probabilities from the host/symbiont and symbiont/gene DTL reconciliation parameters instead of using an additional parameter (described in the previous sections).

The faster Sequential heuristic may not be as robust as the Monte Carlo one. Li and Bansal [27] present an example where the sequential approach cannot propose a solution at all, in a parsimony model where inter horizontal gene transfer are forbidden. In figure 3 we present another illustration, with this time an emphasis on the “not continuous” aspect of the Sequential heuristic in regard to the host and symbiont reconciliation events rates.

**Figure 3:**
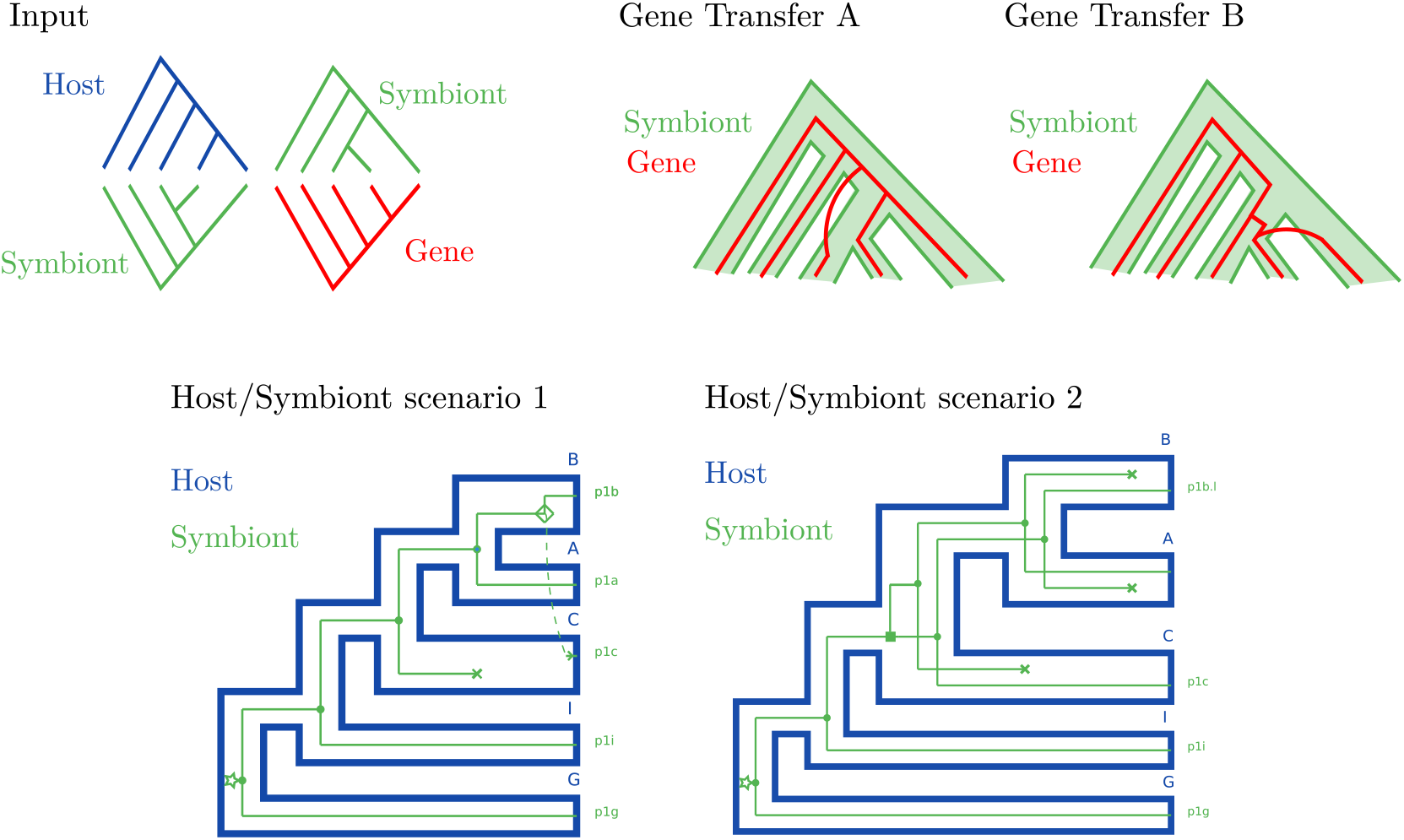
An example of input where the Sequential heuristic is less robust than the Monte Carlo one. We compare the support for two gene transfer scenarios, scenario A and B. There are two main possible host/symbiont reconciliation scenarios, scenario 1 and scenario 2. In scenario 1, gene transfer A is more likely, and in scenario 2, gene transfer B is more likely (both gene transfers involve ghost species, whatever the scenario. The support for both gene transfers for the Sequential and Monte Carlo heuristics are presented in table 1.

**Table 1:** Comparison of the support for the two gene transfer scenarios in the example presented in figure 3. Column 1 contains the method: Monte Carlo (Section 3.2), Sequential (Section 3.4) and 2-level (which consists in reconciling *G* with *S* without information form *H*). Then columns 2 and 3 contain the support of gene transfers, respectively *A* and *B* (in reference to Figure 3), according to reconciliation scenario 1 or 2, obtained with different transfer probabilities.

| Heuristic | Gene transfer A | Gene transfer B |
| --- | --- | --- |
| Host Symbiont rates **T 0.006 D 0.1 L 0.1** |  |  |
| Monte Carlo | 0.43 | 0.27 |
| Sequential | 0.90 | < 0.05 |
| 2-level | 0.18 | 0.21 |
| Host Symbiont rates **T 0.005 D 0.1 L 0.1** |  |  |
| Monte Carlo | 0.35 | 0.33 |
| Sequential | < 0.05 | 0.49 |
| 2-level | 0.19 | 0.23 |

A small change in the transfer rate of the host and symbiont makes a big difference for the gene and symbiont reconciliation with the Sequential heuristic, but a small one for the Monte Carlo one, see the results in table 1.

### 3.6 Time complexity and tractability

We denote *h*, *s*, *g* the number of nodes of the host, symbiont, and gene trees respectively. It has been demonstrated that 2-level parsimony DTL reconciliations can be computed in quadratic time [6] if all transfers have the same probability.

In our implementation sampling one host symbiont reconciliation scenario (line 8 in Algorithm 1) is done in cubic time *O*(*hs*^2^) complexity because we parse the transfer sum from equation 1.

Computing the gene transfer probabilities between all couples of symbiont nodes (section 3.3) is done with a dynamic programming similar to the one for reconciliation in *O*(*hs*), presented in equation 5. A final sum (equation 4) over all hosts of the considered symbionts in *O*(*h*^2^*s*^2^), in the reasonable case where the number of symbiont nodes per host nodes (in the reconciliation scenario) is below a constant k, yields *O*(*h*^2^*k*^2^ + *hs*) for this part.

Finally the host aware gene/symbiont reconciliation (line 15) differs with classic 2-level reconciliation in that transfer rates depend on the donor-receiver couple. In consequence we cannot use the efficient computation trick used for uniform rates [6, 56], that enable to compute equation 1 without computing for each couple of gene and symbiont subtrees the transfer sum. Here for each couple of gene and symbiont subtrees, we must explicitly consider transfers toward all symbiont nodes, yielding a cubic complexity of *O*(*s*^2^*g*) for host aware symbiont/gene reconciliation.

This leads to a total complexity of *O*(*N* (*hs* + *h*^2^*k*^2^ + *s*^2^*g*)) where *k* is a bound on the number of symbionts per host in the sampled reconciliations (*s* in the worst case), and *N* is the number of samples in the Monte Carlo approach.

The datasets presented give a good idea of the size of the data we can consider with this new method. We here give the computation time for the Sequential heuristic. Computation on the Cinara aphid dataset, with a size of 25 leaves for the symbiont tree, 9 leaves for the host, and 13 gene families takes about 3 minutes on a single laptop core, including the rate estimation steps. This is a dataset on which it would be possible to use the Monte Carlo approach. The pylori dataset is larger, the symbiont has 119 leaves, the host 7 leaves, and there are 1034 gene families, of which 322 have 119 leaves. Reconciliation, with fixed rates (without rate estimation) took just under a day using 8 cores.

### 3.7 Symbiont tree inference

In case the symbiont tree is unknown, we devised an option to infer the symbiont tree by amalgamation [12, 56] of universal unicopy gene trees, guided by the host tree.

Clade prior probabilities are computed from universal unicopy gene trees, and dynamic programming is used to compute the likelihood. A symbiont tree is sampled in the backtracking phase at the same time as the host/symbiont reconciliation scenario. This amalgamation is also implemented for the symbiont/gene part, to account for gene tree being unrooted, and to be able to include uncertainty in gene tree topology, just like in 2-level reconciliations[21, 56].

### 3.8 Rates estimation and likelihood comparison

In our model, the data is the gene trees, and the free parameters are the three DTL probabilities of the symbiont/gene reconciliation. We consider the host/symbiont DTL parameters as fixed, *i.e.* estimated without knowing the data. This makes it possible to compare, based on the likelihood, our approach and a 2-level one (symbiont/gene reconciliation, unaware of the host), because they have the same free parameters, and because they both define a probability distribution on the same space of gene trees associated to the symbiont tree.

In practice we estimate the host/symbiont DTL parameters, as done in ALE [57], with an expectation maximization method, and then fix these parameters. Then we run the Monte Carlo or sequential approach multiple times to estimate rates for the symbiont/gene reconciliation with the same expectation maximisation method.

### 3.9 Output format and solution visualization

Our implementation can output a sample of full scenarios, both for symbiont/genes and the corresponding host/symbiont reconciliations. The scenarios are given in RecPhyloXML, a common standard for reconciliation output endorsed by a significant part of the gene/species reconciliation community [18]. The scenarios can be visualised using Thirdkind^1^ [41], a reconciliation viewer that handles 3-level reconciliations. We also output event frequencies based on the reconciliation scenario sampling. Indeed we sample a number (100 by default) of symbiont/gene reconciliations and observe the frequency of each event in these replicas, thus obtaining an estimate of the posterior probability of events. It is this result that we use to evaluate the ability of our method to infer specific events, such as receptors and donors of horizontal symbiont transfers, which we compare to simulated scenarios or previously proposed scenarios on aphids Cinara.

## 4 Experimental results

### 4.1 Simulated datasets

#### 4.1.1 Description of the simulation process

Our probabilistic model can be used for simulation, however in order to test our method, we chose to use an exterior simulation framework. We used the available software Sagephy developed by Kundu and Bansal [24]. Sagephy generates three embedded trees and allows replacing transfers on top of additive ones. We used the parameters proposed by the same team in another article [23], as representative of small (D 0.133, T 0.266, L 0.266), medium (D 0.3, T 0.6, L 0.6) and high (D 0.6, T 1.2, L 1.2) transfer rates, without replacing transfers. The software enables to specify an inter transfer rate, corresponding to the probability for a gene transfer to be between symbionts hosted by different hosts (”inter” transfer). When a horizontal transfer is chosen during generation of the gene tree (inside a symbiont tree and knowing a host/symbiont reconciliation), the transfer is chosen to be an inter host one with the inter transfer rate. So an inter transfer rate of 0 corresponds to our inference model of only intra transfer, and of 1 corresponds to a case where transfers are only between symbionts in separate hosts.

We constructed two simulated datasets, one with a combination of the different rates for the DTL parameters, and one with only medium rates but with different inter transfer rates. For the first dataset, we used all 9 combinations of small, medium and high rates for the symbiont generation and the gene generation, with only intra host gene transfer (*i.e.* an inter transfer rate of zero). For the second dataset, we used only medium rates for both symbiont and genes generation, but we used 6 inter transfer rates going from 0 to 1.

For both datasets, and for each set of rates, we generated 50 instances consisting of 1 host tree with 100 leaves, 1 symbiont tree and 5 gene trees, each generated in the pruned version of the other trees (branches that do not reach present are pruned before the generation of the next tree). We then selected host leaves with a probability of 0.08 to simulate unexhaustive sampling, resulting in host trees with an average size of 8 leaves. We thus simulate extinct lineages, and even with a simulation inter transfer rate of 0, some gene transfers will be inter. This ended up to 399 instances for the first dataset and 226 instances for the second one, and at least 29 instances of 5 genes for each set of parameters.

We compared the results from three approaches. (1) The “2-level” heuristic which is a 2-level reconciliation between the gene and symbiont trees, ignorant of the host tree. (2) The “Sequential” heuristic, which consists in computing the most likely host/symbiont DTL reconciliation and doing the symbiont/gene reconciliation, given that host/symbiont reconciliation. (3) The full 3-level “Monte Carlo” method, summing the results of the gene reconciliations over 50 sampled host/symbiont reconciliation scenarios. We let our approaches estimate evolutionary rates.

We measured first the capacity of the three methods to infer the correct symbiont donor and recipient of gene transfers (with precision and recall), and second, the likelihood they attribute to symbiont/gene cophylogeny. Identifying the exact donor and recipient of simulated transfers is usually considered a hard task for reconciliation algorithms. Usually reconciliation studies are not evaluated with this strong criterion [37], but with the inference of ancestral characters [60], the number of transfers [54], the ability to infer better trees [8], or the ability to map the correct event type to each gene node [23]. We chose to look at the capacity to infer specific transfers because we feel that it is in this task that our model has the capacity to show its utility. It can infer more precise gene transfers because transfers are constrained by additional elements compared to other methods.

Our probabilistic reconciliation approaches output estimates of the posterior probabilities of evolutionary events, so we used these probabilities as weights for our precision and recall definition in Figure 4 for the detection of horizontal gene transfer donor and receiver symbionts. Denoting by *L_t,sim_* the list of simulated transfers and *L_t,obs_* the list of observed transfers, and *P_obs_*(*T*) the estimation of our approach for the probability of transfer *T*.

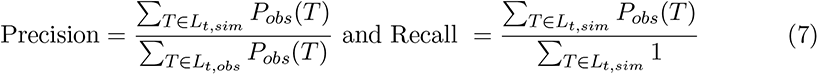

**Figure 4:**
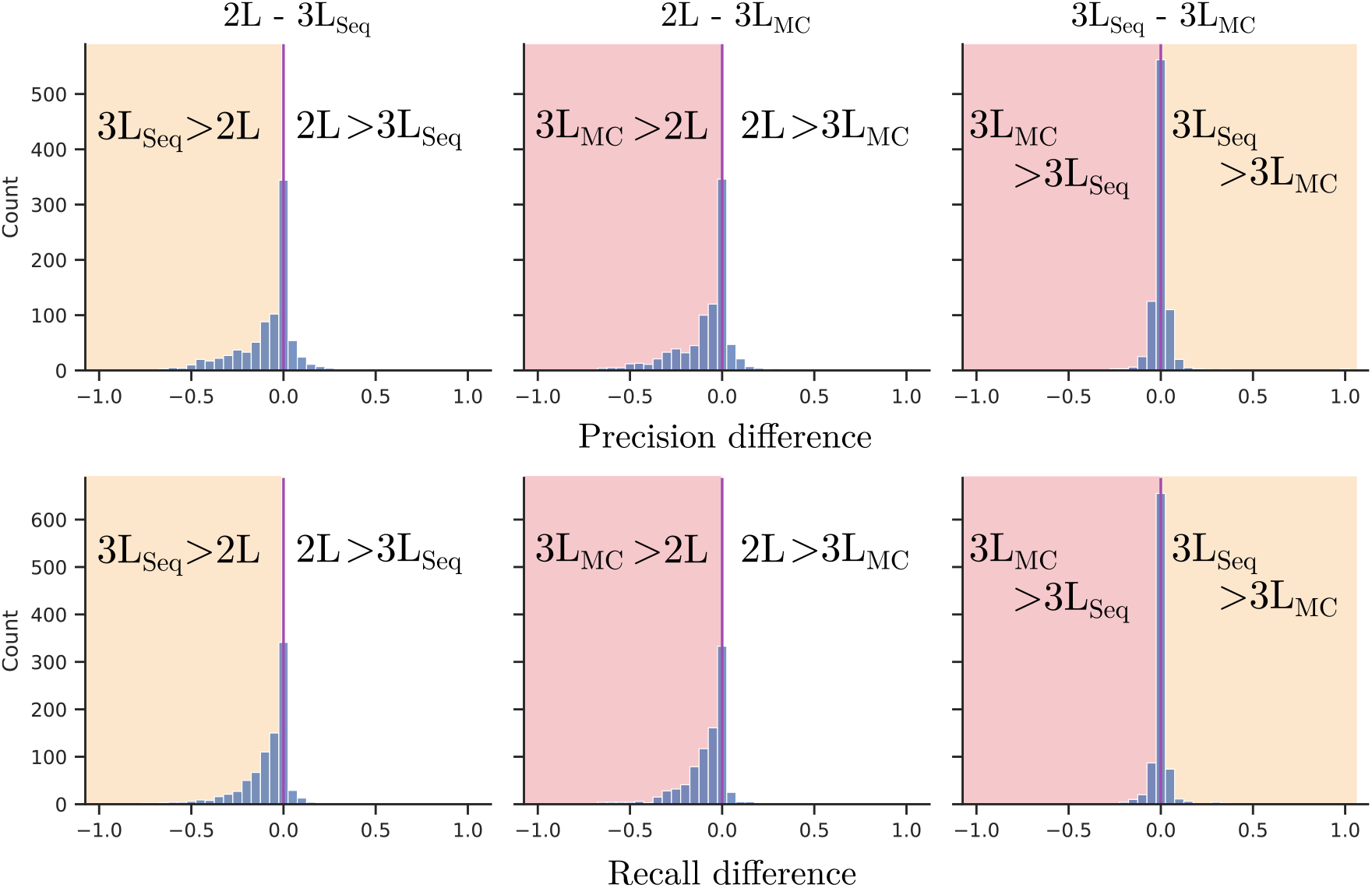
Distribution of differences of precision and recall on the inference of horizontal gene transfers for all combinations of two approaches: 2-level (2L), 3-level with the Monte Carlo heuristic (3L_MC_) and 3-level with the Sequential heuristic (3L_Seq_), centered on 0, and for all 874 gene families of the 3-level simulation, with no inter host gene transfer, that undergo at least one transfer.

#### 4.1.2 The 3-level method infers more true transfers than the 2-level method

Overall the Monte Carlo and sequential approaches give similar results on these simulated datasets, and better results (in particular for recall and to a lesser extent for precision) than the 2-level approach (Figure 4). In most cases, the faster Sequential heuristic can advantageously replace the Monte Carlo one because they have the same recall and precision. In a few case, that might be the more interesting ones, the Monte Carlo has a slight advantage, and though it is more computationally costly, it is also theoretically more robust.

In addition we measured the time consistency of reconciliation scenarios in the 2-level and 3-level inferences. Indeed, we have already remarked that we work in an undated framework, and in consequence transfers might be incompatible [13]. For each simulation condition we listed all inferred transfers and checked compatibility. For 2- level reconciliations, 35% of the conditions lead to time incompatibilities, this same measure dropping to 15% if 3-level reconciliations were performed.

#### 4.1.3 A host-symbiont co-evolution test

The reconciliation likelihood difference between 3-level inference and 2-level inference is a marker of host-symbiont co-evolution. Indeed, Figure 5 (A) shows that when the simulation model is less dependent from the host phylogeny (inter transfer rates of 0.6, 0.8 and 1.0), the likelihood difference between the 2-level and 3-level inference methods are mostly in favor of the 2-level. It happens for almost all instances in the simulation dataset with with no intra transfers (inter transfer rates of 1.0), the farthest one from the model behind our heuristic that privileges intra transfers. For all these instances a preference for 2-level reconciliation (according to the likelihood) is more likely when few transfers are inferred (we sum over 1 to 5 gene families generated for each host and symbiont instance). This is a sign of the precision of the method to not classify 2-level instances as 3-level ones.

**Figure 5:**
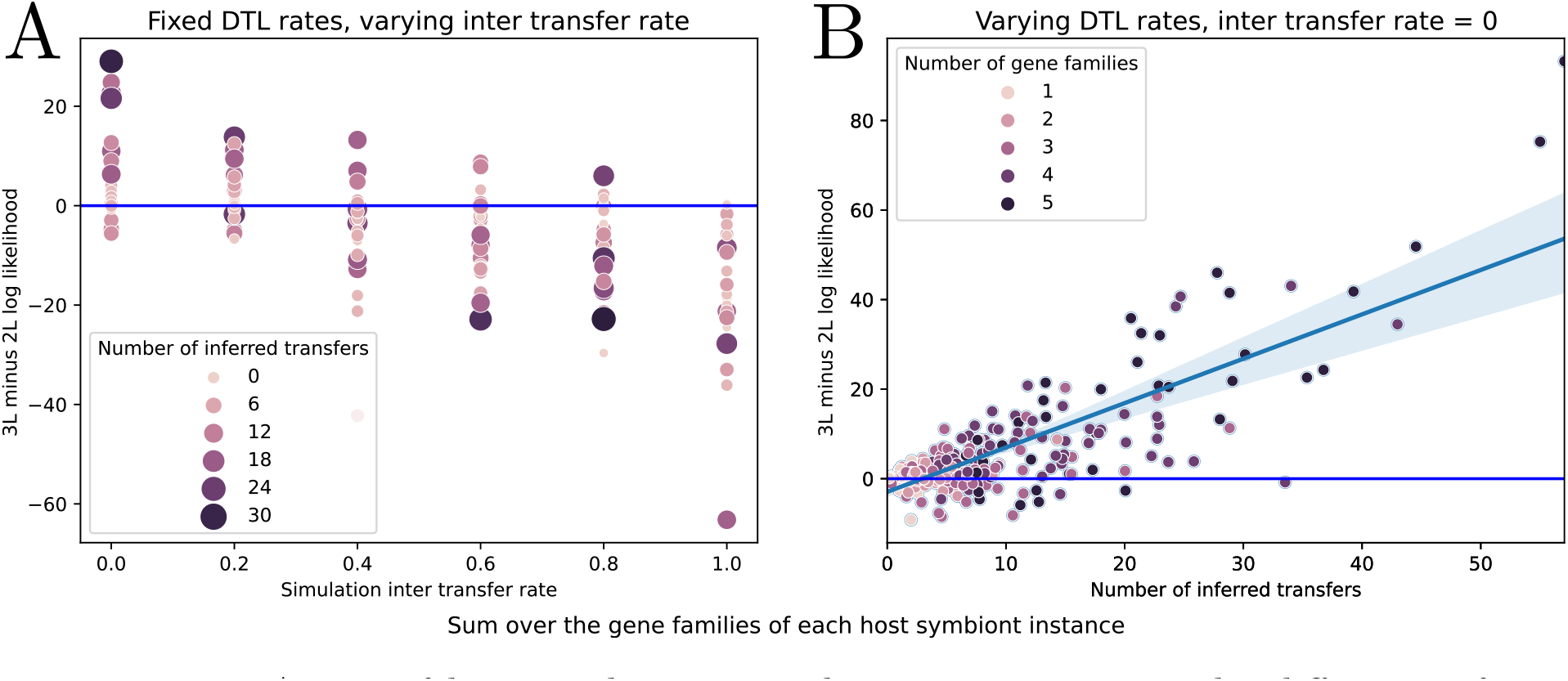
A test of host symbiont co-evolution. We measure the difference of likelihood between the 3-level model and the 2-level model, using the estimation of these likelihoods provided by our “2-level” and “3-level Sequential” heuristics, in order to differentiate instances where gene trees are generated in a 3-level host/symbiont/gene model or in a 2-level symbiont/gene model. Each instance is composed of a host tree, a symbiont tree, and 1 to 5 gene families. For one instance we sum the differences over all gene families. (A) Sensitivity of the likelihood difference to the value of the inter host gene transfer probability in Sagephy. As expected, the more an inter transfer rate is probable (independent from the host phylogeny), the less we detect host-symbiont co-evolution with the likelihood difference measure. Colors indicate the number of inferred transfers. (B) Sensitivity of the likelihood difference to the number of inferred transfers (dataset with only intra transfers). Colors depict the number of gene families considered in the host and symbiont instance. Because transfers carry the co-evolution signal, the sensitivity of the method increases with the number of transfers, which are higher if we increase the number of gene families.

In a model with only intra transfers (inter transfer rate of 0), we have a very good recall for the detection of the 3-level model, almost all only intra transfer instances are classified as 3-level as they should be. A more detailed exam of this recall is presented in Figure 5 (B) with the first simulated dataset, with only intra transfer and varying DTL parameters.

Figure 5 (B) shows the likelihood difference when only intra transfers occur in the simulations. We see that when the number of transfers is higher, the likelihood difference better reflects the mode of simulation. In practice a way to increase the number of transfers is to increase the number of gene families considered.

### 4.2 Precise identification of a gene transfer in enterobacteria symbiotic of *Cinara* aphids

A recent study on Cinara aphids enterobacteria systems [28] identified one host switch and two horizontal gene transfers, one intra-host from *Erwinina* to *Hamiltonella* and one inter-host from *Sodalis* to *Erwinia*. The genes transferred (thi) and some others (bioa,d,b) were first inherited through gene transfers, probably from *Sodalis* related symbionts. Moreover, those genes transferred are part of functions to complement the lack in the sap-feeding host nutrition. It seems that a new endosymbiont acquires the genes of another one to sustain the host. This exemplifies a case where a symbiont gene can co-evolve with the symbiont host, more than with the symbiont itself. We reproduced this scenario in Figure 6 (A), and a representative gene tree witnessing the transfers is reproduced in Figure 6 (B).

**Figure 6:**
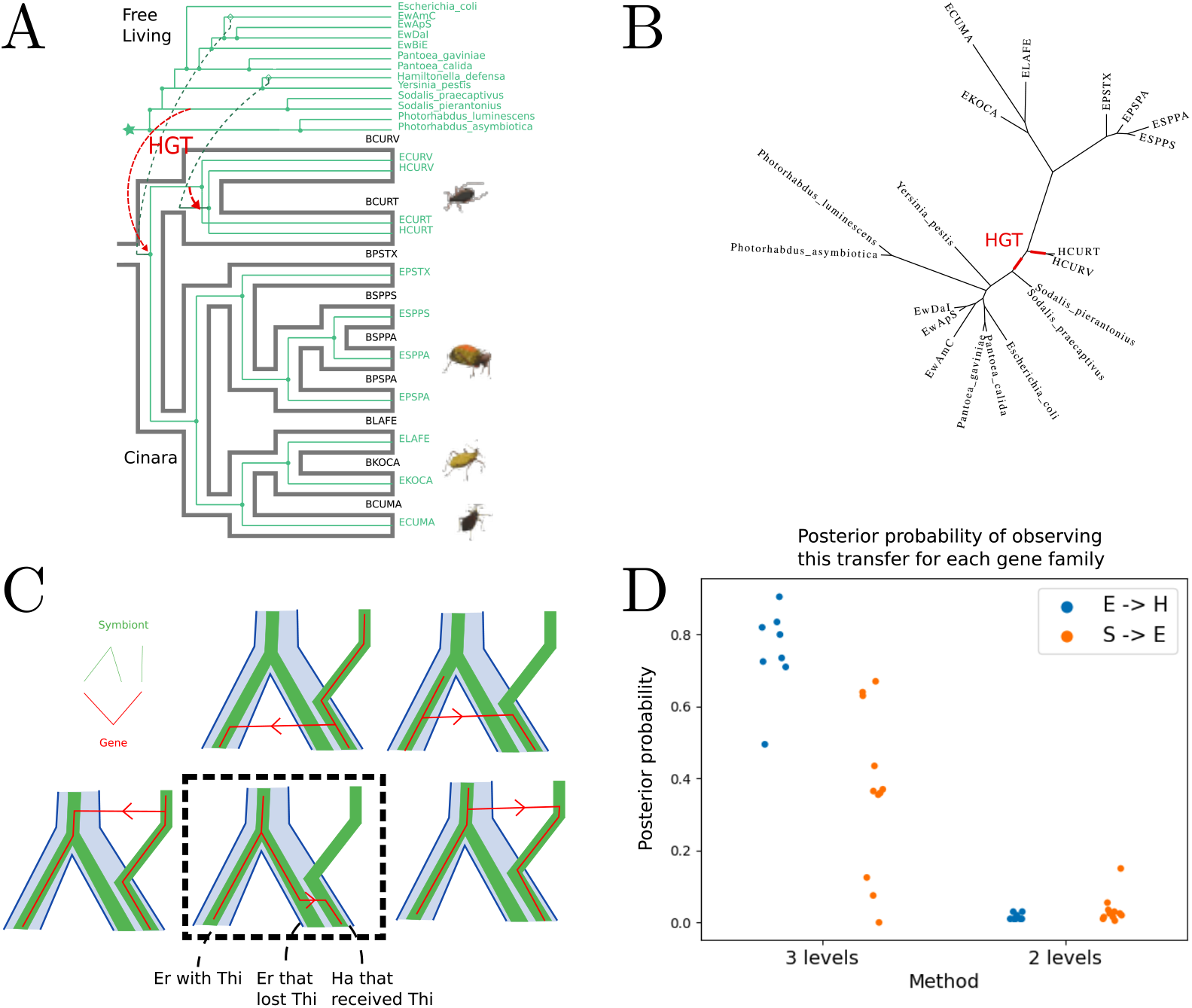
The evolution of *Cinara* and their enterobacteria symbionts. (A) The evolutionary scenario identified by Manzano-Marín et al. [28]. The reconciliation of the hosts (*Cinara* aphids) and symbionts (bacteria) are depicted along with the position of the horizontal gene transfers (in red). (B) Phylogenetic tree of one gene with the position of the two transfers. (C) Theoretical explanation of the difference between the results of the 2-level and 3-level reconciliation methods. The two top reconciliations are a bit more likely in a 2-level framework, as they require a single transfer while the bottom ones require a transfer and a loss, but one of the bottom one (with the dotted square) is better in a 3-level model, as it allows an intra-host transfer. (D) Support (a posteriori probability of the transfer, computed from its observed frequency in the reconciliation sample) for the identified HGTs, from *Erwinia* to *Hamiltonella*, and from *Sodalis* to *Erwinia*, for 3-level and 2-level reconciliations.

Gene trees including *Cinara* endosymbionts and other enterobacteria species were available from the supplementary material made available by Manzano-Marín et al. [28]. *Cinara* and their endosymbionts phylogenies show exact correspondences on the studied period. We kept all enterobacteria associated to a *Cinara* aphid (of *Erwinia* and *Hamiltonella* genus), and chose a representative subset of the other enterobacteria present in the gene trees, notably containing *Sodalis* species, closest identified parent to one of the transferred genes, and other *Erwinia* and *Hamiltonella* genus species. We used the phylogeny proposed in Annotree for these species [32], to complement the *Cinara* aphids symbionts phylogeny proposed in [28]. We used our 3-level reconciliation on the host tree and symbiont tree, using the possibility of our method to take into account these “free living” bacteria. As the host and symbiont (apart from the free living) are identical, we used the sequential heuristic.

We tested the capacity of the 3-level method compared to a 2-level one to detect the gene transfers identified by Manzano-Marín et al. [28]. The intra transfer from *Erwinina* to *Hamiltonella* is retrieved in around 80 percent of the scenarios sampled by the 3-level method, and both are better retrieved than in the method that does not take the host into account (Figure 6 (D)). A theoretical explanation using a toy example is given in Figure 6 (C). An alternative transfer, in the other direction, from *Hamiltonella* to *Erwinia* is slightly more likely but the configuration of the host evolution supports the intra transfer.

This exemplifies how multi-scale dependencies can only be captured by 3-level models.

### 4.3 *Helicobacter pylori* genes as documents for human migrations

*Helicobacter pylori* is a bacterial symbiont of a significant proportion of humans, which has been supposed to be a marker of human migrations across the Earth [1]. Bacterial strains have been divided in different populations corresponding to geographical areas (Africa 1, Africa 2, Asia 2, East Asia, North East Africa, Europe) [59, 30].

The supposed coevolving complex made by humans, bacterial symbiont and their genes makes it an accessible system for the host/symbiont/gene reconciliation method. In particular gene transfers should be more probable between *Helicobacter* strains if they are hosted by a same human population.

We collected available current strains of *H. pylori* from the NCBI which have a genetic population assigned by MLST allelic profile [2, 22]. A phylogenetic tree was built based on the concatenation of universal-unicopy genes (322 gene families), and a sample of 113 strains representing the diversity of H. pylori in the old world (excluding strains from the Americas) was obtained using Treemmer [31]. Then, 6 non pylori strains were added (*H. hepaticus, H. acinonychis, H. canadensis, H felis, H. bizzozeronii, H. cetorum*), as external groups.

In this study we considered the 1034 gene families, including 322 universal unicopy families, that displayed strains from the external groups and from at least 3 continents. We then considered four different population trees (host trees) containing the geographical areas as leaves, coherent with the scientific literature [59, 30]. 322 universal unicopy gene trees were used, and the strain (symbiont) tree was amalgamated from gene trees with the population trees as a guide (see subsection 3.7). As strains were much more numerous than populations, and subject to a more complex diversification than DTL events, we allowed an additional event, named I, that consists in a duplication followed by a speciation and loss of one of the copies, with a specific rate, inferior to the combination of these three events. This event allows a strain to be present in a population and one of its descendants, and is used as one of the default events in biogeography frameworks [45].

We then applied our sequential approach and compared the likelihood of the gene-/strains aware of the host reconciliation to compare the population trees. The results are depicted in Figure 7 (A). The likelihood of the systems according to the population tree is reported, divided into two components: the likelihood of the population/strain comparison, and the likelihood of the gene/strain aware of the population comparison. The population tree on the left column is the most likely given the model, the method and the used data. Assessing the robustness of the result would require a sensibility study which is out of the scope of this contribution.

**Figure 7:**
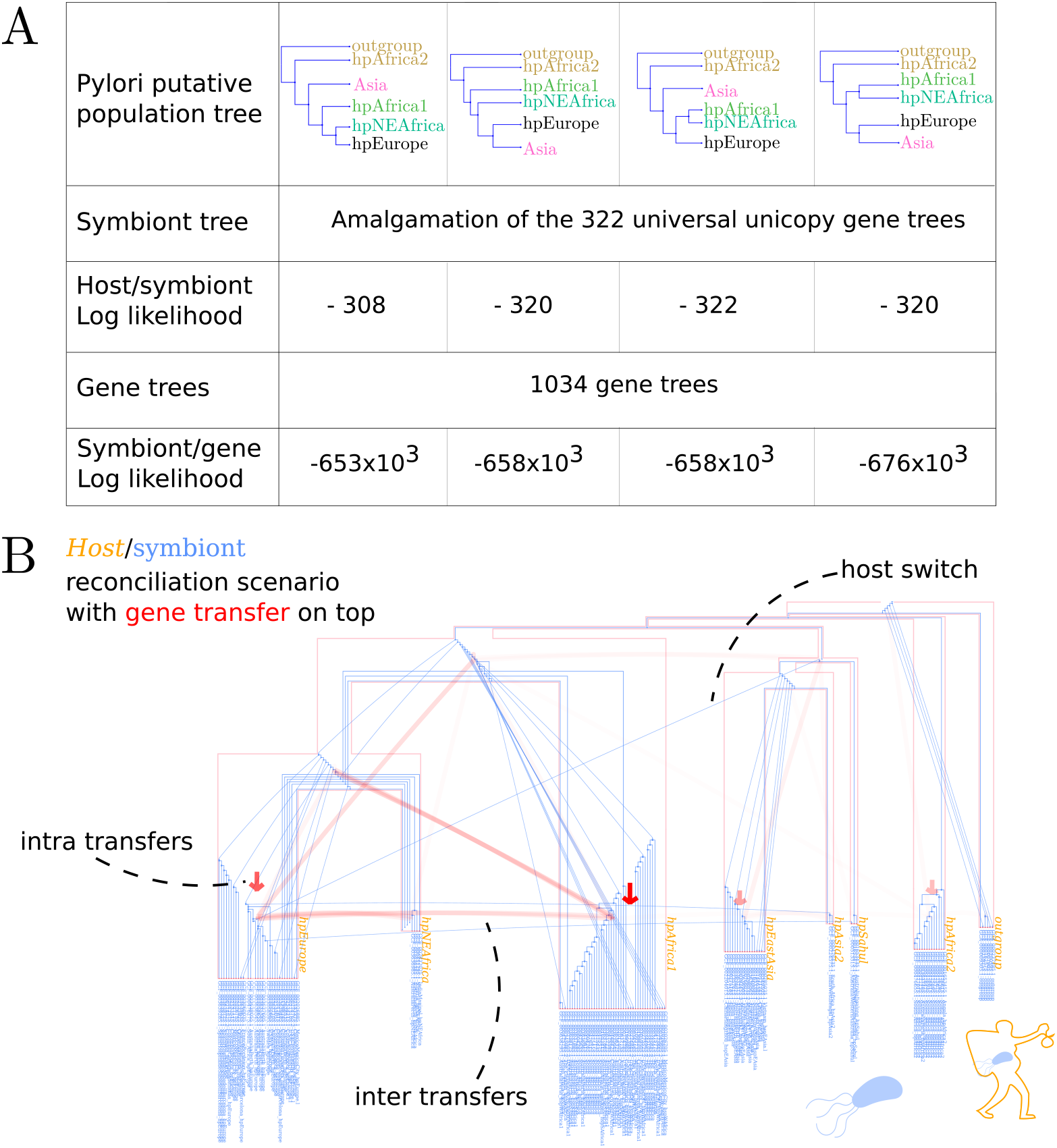
Co-evolution of human populations and *Helicobacter pylori*. (A) Log likelihood of the different population trees. (B) The representation with Third-Kind [41] of one possible reconciliation scenario of *Helicobacter pylori* strain tree and the population tree maximizing likelihood. Aggregated gene transfers are depicted on top of the DTL reconciliation, with the opacity corresponding to the number of times the transfers were seen across the 1034 gene families.

Figure 7 (B) is an illustration of a reconciliation scenario for the maximum likelihood host tree with Thirdkind [41]. We see the host tree and the amalgamated strain tree reconciled (I events are represented as transfers from a parent node to one of its child). On top of these two embedded trees red lines represent the aggregation of gene transfers depending on the host of the donor and receiver strains. The opacity of the transfer lines are proportional to the number of times a certain kind of transfer is observed across the 1034 gene families in one sampled scenario.

## 5 Discussion

In a review on horizontal gene transfer in host symbiont systems [61] the authors highlight the need of plurality of evidence to robustly assess the existence of transfers. Evidence can be of multiple types, gene trees, donor receiver ecology, or host symbiont association. We provide a framework were these multiple evidence can be gathered, and the proof of concept that it can work, on *Cinara* aphids and their enterobacteria. Our method uses a probabilistic framework that enables rate estimation, tree inference, tree comparison and model comparison. We also introduced a method to compute the inter transfer rate from the intra transfer one and the modeling of ghost lineages in the host symbiont reconciliation. We introduced a Monte Carlo approach that enables to estimate event probabilities and likelihood, by sampling through multiple host symbiont scenarios in a double DTL model. Implementation is available on GitHub https://github.com/hmenet/TALE.

While our intuition is that the Monte Carlo approach is more robust than the sequential one, notably in cases where gene events happen around uncertain host symbiont reconciliation nodes, our evaluation on simulated data did not show a big difference in most cases. We think that in biological data, we can expect more interaction between the events of the host symbiont reconciliation and the ones of the gene symbiont one, which are independent in our simulation. Developing new simulation frameworks that can model such dependencies, for instance by increasing the loss rates when multiple genes or symbionts are present, or using a functional approach to the evolution of genes, could be important to the understanding of these multi-level models.

The ability of our inference methods to be used for model comparison seems promising. We saw that with an increasing number of gene families we could increase our confidence in the answer. However the different gene families must contain a part of independent information, as is the case in the simulation where all families dependence are completely in the host and symbiont trees. For instance in the *Cinara* aphids dataset, the genes considered are mostly similar, and do not really make the number of independent transfers increase, and with only one intra transfer, that necessitates an additional loss to occur, the 2-level model displays a better likelihood than the 3-level. If more independent transfers were present, we can suppose that some of them might not necessitate such a loss and the test would favor a host symbiont co-evolution.

All these features deserve further tests to know their domain of validity and to draw biological conclusions. In particular, the inference of the symbiont tree, with the use of amalgamation, from an input distribution of universal unicopy gene tree would deserve to be tested against other standard methods as concatenate or species tree reconstruction with 2-level reconciliation model as it is implemented in SpeciesRax [35].

An interesting future direction in this line would be to construct, instead of a symbiont tree, compartment trees, which would depict the evolution of inter-dependent genes that are not necessarily in the same species.

A comparison of the inference method to similar ones [52, 26, 36] could also be undertaken. However in an host/symbiont/gene framework, horizontal transfer in the host/symbiont reconciliation are crucial, and only the model of Stolzer et al. [52] takes these events into account. Moreover the sequential heuristic is simply a rewriting of this model in a probabilistic framework.

More generally, the model is not bound to host/symbiont/gene systems, but any set of three nested inter-dependent entities can be studied with it: species/gene/protein domain as it was done in previous studies [52, 25, 36], or geography/species/gene, and so on. As the scales of biological observation are probably infinite, so are the combination of three nested scales.

Examples presented in this article show the possibilities of the method, but still derive no biologically significant breakthrough. However the necessity of such a method, detecting multilevel co-evolution, could arise with the more and more numerous studied biological systems that fit into this multi-scale cophylogeny framework, notably with an increasing interest for hologenomics [3].

## Supporting information

Supp_figures_cinara_thirdkind

## 6 Acknowledgements

This work was performed using the computing facilities of the CC LBBE/PRABI.

## Footnotes

1 https://crates.io/crates/thirdkind

