## Supplementary material for "Host-symbiont-gene phylogenetic reconciliation": Supp_figures_cinara_thirdkind

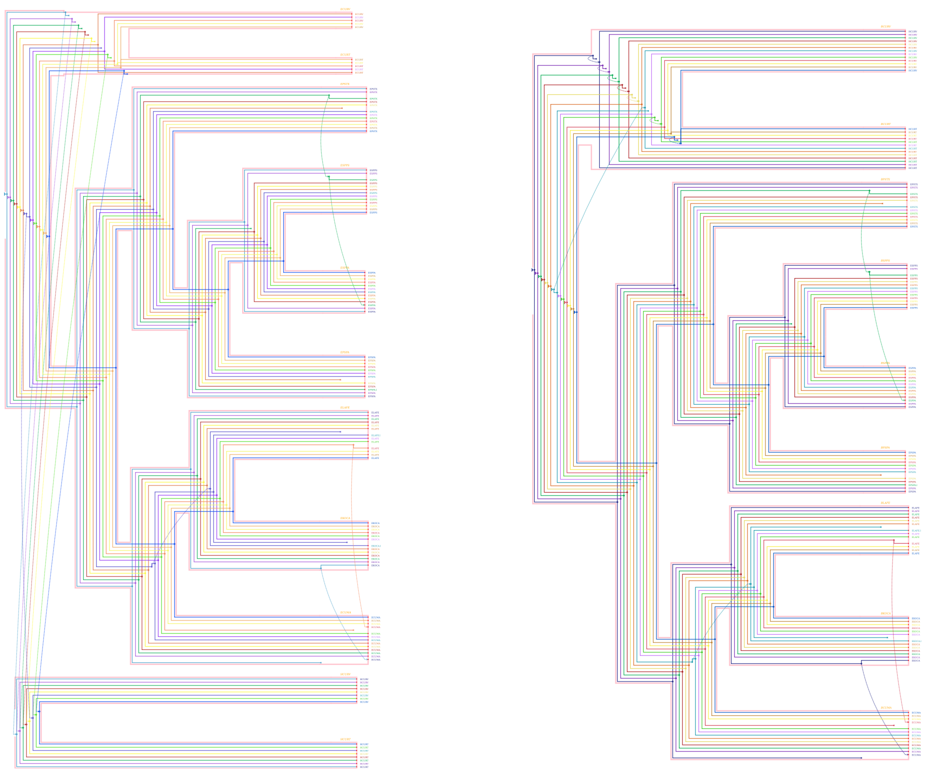

Figure 1: Views of *Cinara aphids*, endosymbionts and genes. Left the symbiont and genes reconciliation (only for the symbionts in the host), and right, the mapped 3 view of symbiont's genes inside the host, we see the genes follow almost exactly the host while there are important transfers between symbionts, with a 3-level approach.

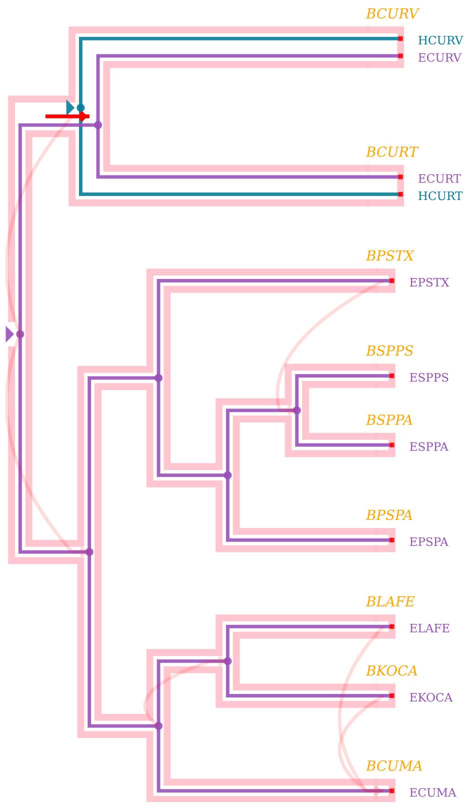


Figure 2: Results of a 3-level reconciliation. Host and symbiont scenario between the *Cinara* host, given by its *Buchnera* symbiont, and the *Erwinia* and *Hamiltonella* endosymbionts. Horizontal gene transfers are represented on top of this reconciliation, with opacity depending on the number of times the transfer is seen across the different gene families. The transfer between Erwinia and Hamiltonella inside their common host is represented as a straight line, depicting a transfer inside a host. Other transfers, specific to single gene families, are also inferred. The figure was generated using Thirdkind, a viewer we present in section.

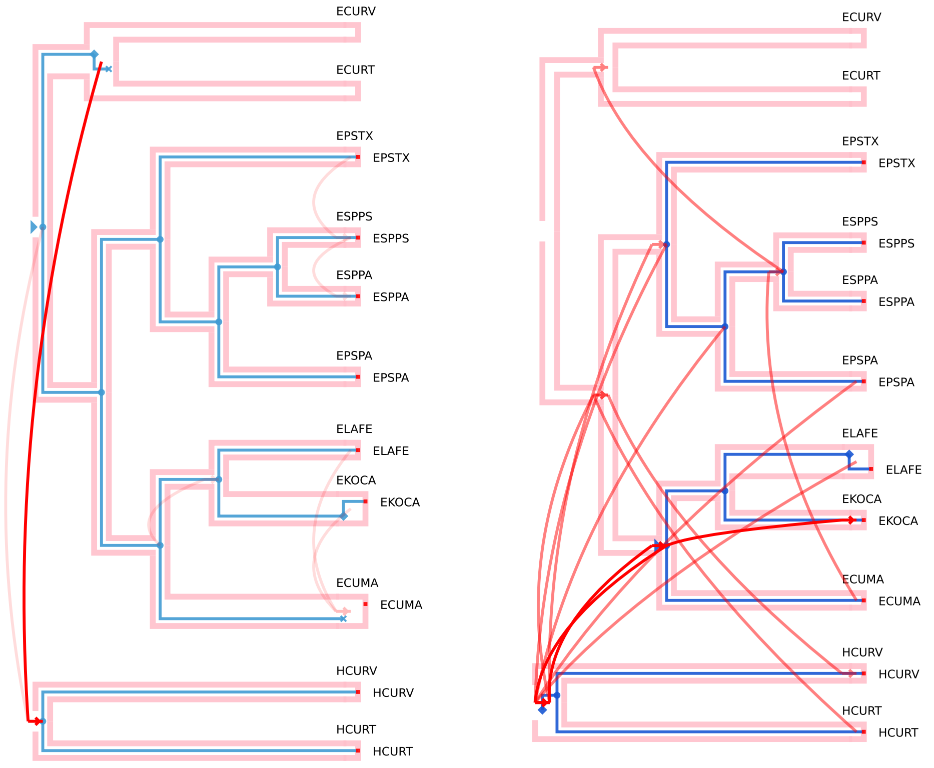


3-level reconciliation 2 level reconciliation

Figure 3: Symbiont and gene reconciliation with the 3-level and 2-level models. One gene family is represented, and, as in figure 2 are aggregated over all gene families. The figure was generated using Thirdkind, a viewer we present in section.
